# A novel highly divergent enteric calicivirus in a bovine calf, India

**DOI:** 10.1101/2023.11.20.567882

**Authors:** Naveen Kumar, Rahul Kaushik, Pragya Yadav, Shubhankar Sircar, Anita Shete-Aich, Ashutosh Singh, Yashpal Singh Malik

## Abstract

In 2015, a novel highly divergent bovine calicivirus was detected in an Indian calf with enteritis. Phylogenetic analysis linked it to the *Nebovirus* with only 38.5% sequence identities, emphasizing the need for separate taxonomic classification. Furthermore, PCR screening detected these unique caliciviruses widely in India’s northern states.

## Text

*Caliciviridae* is a family of non-enveloped viruses distinguished by their single-stranded, positive-sense RNA genomes of 7.4–8.3 kb and characteristic “star-like” appearance under electron microscopy. *Caliciviridae* members cause a wide spectrum of illness in a broad range of animals and humans (1). It currently encompasses eleven genera, seven of which (*Lagovirus, Norovirus, Nebovirus, Recovirus, Sapovirus, Valovirus*, and *Vesivirus*) infect a wide variety of mammals, two (*Bavovirus* and *Nacovirus*) infect birds, and the remaining two (*Minovirus* and *Salovirus*) infect fish (1).

Enteritis in bovines has been linked to caliciviruses from two genera, *Nebovirus* and *Norovirus* (2-5). However, *Vesivirus* infections have been found in bovines without causing enteritis (6). We present the findings of a bovine enteric calicivirus, which is distantly related to neboviruses, in a calf with enteritis in India.

## The Study

In the month of May of 2015, 14 cross-bred calves less than a year old reared in the temperate high-altitude Himalayan region (Mukteshwar, Uttarakhand, India), were presented with severe diarrhea, weight loss, and dehydration. To identify the causative pathogen, fecal samples (n = 14) were collected aseptically from the infected calves, all of which tested negative for genome of rotavirus group A (RVA) (7), Picobirnavirus (PBV) (8), Bovine Coronavirus (BCoV) (9), and Astrovirus (AstV) (10). However, the RIDASCREEN Norovirus ELISA kit (R-Biopharm, Darmstadt, Germany) detected the possible presence of *Norovirus* in one of the samples, UK-B6. We archived this sample at -80 °C for future use. The next generation sequencing (NGS) was performed directly on the fecal sample at the ICMR-National Institute of Virology, Pune, India and the complete genome sequence data retrieved was submitted to the National Centre for Biotechnology Information (NCBI) (GenBank accession no. MN241817).

The genome size of bovine calicivirus (UK-B6) retrieved from sample was 7,484 nt long, with a 91 nt long 5′ untranslated region (UTR), two open reading frames (ORF), ORF1 of 6,657 nt and ORF2 of 657 nt in length, and an 80 nt long 3′ UTR. ORF1 encoded a large polyprotein of 2,218 amino acids (aa) in length, which included non-structural proteins as well as a 542 aa long VP1 protein (capsid protein) at the 3′ end. ORF2 encoded a protein that was 218 aa long. The ORF1 polyprotein contained well-preserved amino acid motifs that are unique to caliciviruses (Appendix Table 1). Furthermore, the International Committee on Taxonomy of Viruses (ICTV) has established a criterion for demarcation of distinct genera within the *Caliciviridae* family, requiring that members in each genus should have more than 60% amino acid sequence divergence in their complete VP1 protein (1). The VP1 protein of UK-B6 differs from all other caliciviruses by more than 60% in amino acid sequence including the prototypes of the *Nebovirus* (65.8% divergence with Newbury1_DQ013304 and 66.3% with Nebraska (NB)_AY082891), indicating that UK-B6 is a unique highly divergent calicivirus (Table 1). Furthermore, a search for VP1 amino acid similarity using Blastp algorithm (available at the NCBI) revealed that two caliciviruses, Kirklareli virus (YP_009272568) and BoNeV15/2021/CHN (WFD61513), had the highest amino acid identities with UK-B6, with 91.6% and 90.5%, respectively. These caliciviruses constitutes a new group (a possible new genus) differing from all other caliciviruses by >60% amino acid sequence differences in their complete VP1 protein.

**Table 1.** Sequence identities of Indian Bovine Calicivirus (Mukti/India/UK-B6/2015) with all the eleven established genera of the *Caliciviridae* family.

| *Caliciviridae* family | Sequence identities (%) |  |  |  |
| --- | --- | --- | --- | --- |
| (Genus_GenBank accession no._virus name) | Genome_nt | Genome_aa | RdRp_nt | RdRp_aa |
| **Nebovirus**_DQ013304_Newbury_agent_1 | 38.5 | 36.9 | 48.6 | 52.6 |
| **Recovirus**_EU391643_Tulane_virus | 26.8 | 15.9 | 33.4 | 24.3 |
| **Bavovirus**_NC_033081_Chicken_calicivirus_strain_RS/BR/2015 | 30.0 | 18.3 | 38.9 | 31.1 |
| **Minovirus**_KX371097_Fathead_minnow_calicivirus | 26.5 | 13.8 | 32.5 | 19.2 |
| **Nacovirus**_JQ347522_Turkey_calicivirus_isolate_L11043 | 30.2 | 19.5 | 39.3 | 29.7 |
| **Salovirus**_KJ577139_Atlantic_salmon_calicivirus_isolate_Nordland/2011 | 27.8 | 14.0 | 34.0 | 20.4 |
| **Sapovirus**_HM002617_Sapovirus_Hu/GI/Sapporo/MT-2010/1982 | 31.1 | 20.8 | 39.7 | 35.0 |
| **Lagovirus**_M67473_Rabbit_hemorrhagic_disease_virus-FRG | 31.2 | 21.3 | 41.5 | 31.6 |
| **Valovirus**_FJ355928_Calicivirus_pig/AB90/CAN | 27.7 | 15.4 | 34.8 | 24.3 |
| **Norovirus**_AF097917.5_Norovirus_Bo/Newbury2/1976/UK | 28.8 | 16.3 | 37.1 | 29.3 |
| **Vesivirus**_U76874.2_Vesicular_exanthema_of_swine_virus_stain_A48 | 28.3 | 19.4 | 40.3 | 31.0 |
| **Vesivirus**_U13992_Feline_calicivirus_CFI/68 | 29.9 | 20.3 | 41.8 | 33.1 |

Furthermore, the complete genome of Calcivirus from UK-B6 diverged significantly from existing member of *Caliciviridae* family’s genera. The whole genome sequence identities with the *Nebovirus* were the highest among all the *Caliciviridae* family genera, namely Newbury1 (38.5% nt, 36.9% aa), whereas it was less than 32% for the other genera (26.8% to 31.2% na, and 14.0% to 21.3% aa) (Appendix Table 2). The RdRp also showed the highest sequence identity to the *Nebovirus* (Newbury1, 48.6% nt, 52.6% aa). Both the 5′ UTR (91 nt; 75 nt in Newbury1, and 74 nt in NB) and the 3′ UTR (80 nt; 77 nt each in Newbury1 and NB) were longer in UK-B6, whereas the ORF1 was 8 aa longer and the ORF 2 was 7 aa shorter in comparison to Newbury1/NB (Figure 1). Typically, *Nebovirus* prototypes have a 1-nucleotide gap between ORF1 and ORF2, whereas UK-B6 had a 1-nucleotide overlap between these two ORFs.

**Figure 1.**
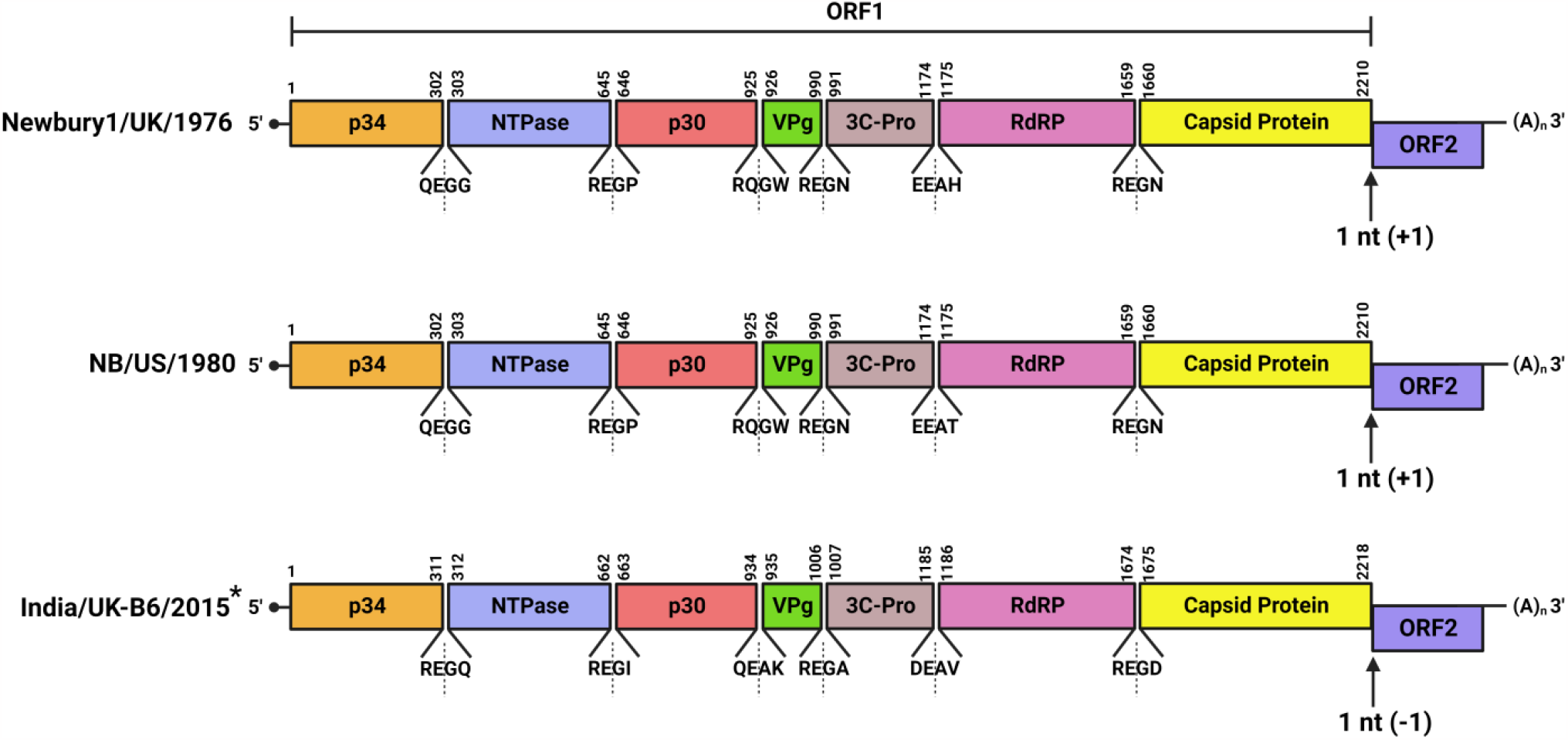
Genome organisation comparison of UK-B6* calicivirus with prototypes of the *Nebovirus* genus, Newbury-1 (DQ013304), and Nebraska (NB) (AY082891). ORF1 encodes a polyprotein which is post-translationally-cleaved into individual non-structural proteins and structural protein (capsid protein). Arrows indicate the putative start codon of the ORF2 at ORF1/ORF2 junction. The putative cleavage sites* on the ORF1-encoded polyprotein of UK-B6 are also shown.

We also compared all the proteins cleaved from ORF1 polyprotein as well as the putative cleavage sites of UK-B6 to Newbury1 and NB, which revealed a longer p34, NTPase, VPg and RdRp, and a shorter p30, 3C-Pro, and capsid protein. UK-B6 calicivirus also had a QEAK cleavage site, which differed from the RQGW cleavage site seen in Newbury1 and NB (Figure 1). Notably, its genome shared the highest whole genome sequence identity with Kirklareli virus (88.6% nt, 94.8% aa), a calicivirus detected in calves with enteritis in Turkey in 2012 (2). Kirklareli virus had similar cleavage sites and proteins cleaved from ORF1 polyprotein to UK-B6 calicivirus.

To dissect the evolutionary dynamics of UK-B6 calicivirus, the genomic region encoding for VP1 protein, a key determinant for phylogenetic separation of distinct genera, was extracted from all eleven genera representative sequences and aligned using MAFFT v.7.475 (11). Based on the Bayesian Information Criterion (BIC), the LG-Gamma substitution model with empirical amino acid frequencies (LG+G+F) was chosen as the best fit amino acid substitution model for the dataset. The phylogenetic trees were then estimated using the Maximum Likelihood (ML) inference and ultrafast bootstrap with 1000 replicates as implemented in IQ tree v2.1.2 (12). UK-B6 calicivirus formed a distinct monophyletic cluster with *Nebovirus* prototypes (Figure 2). Nonetheless, the short branch length of UK-B6 calicivirus suggests an ancestor of *Nebovirus* genus prototypes.

**Figure 2.**
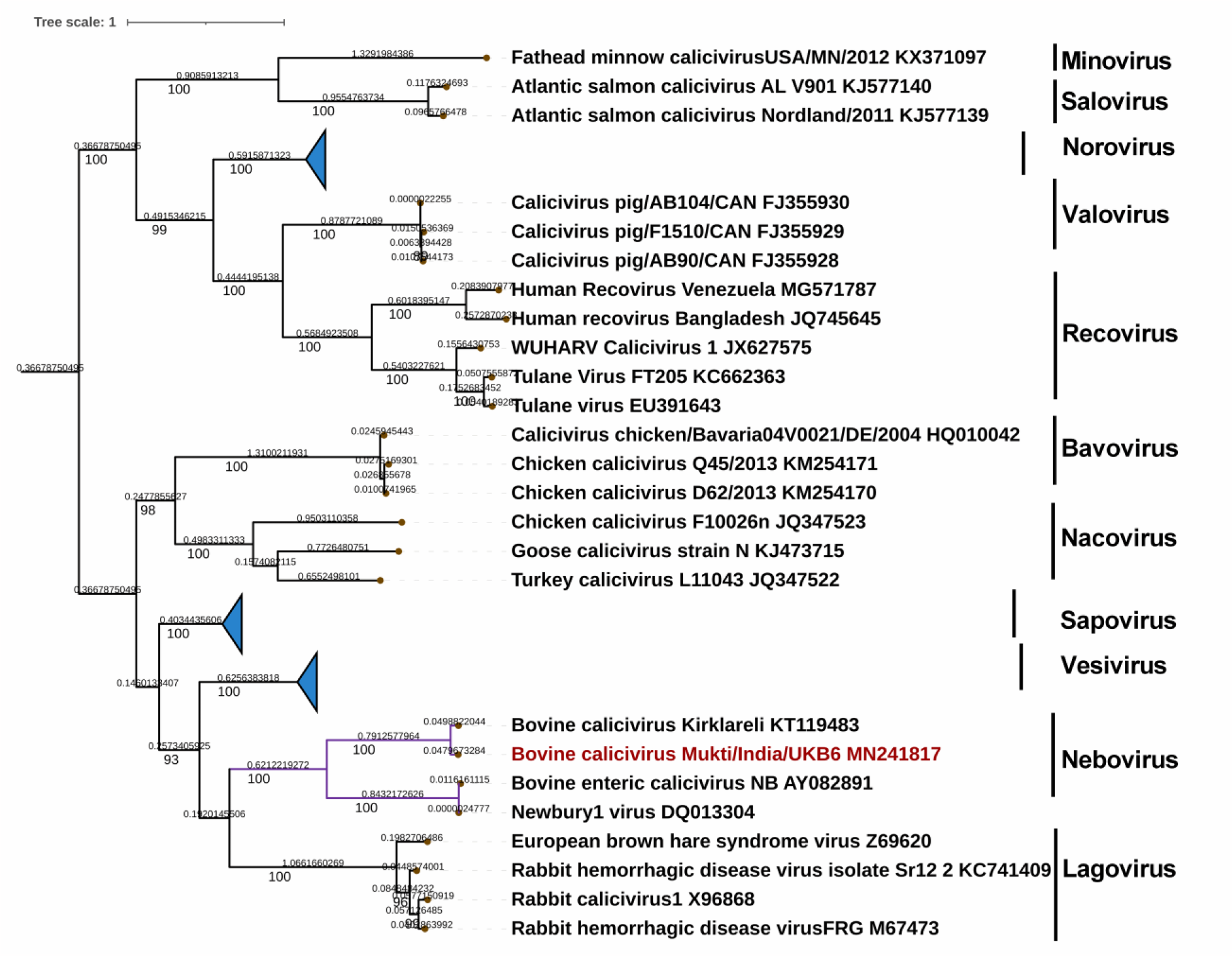
The maximum-likelihood tree depicting the phylogenetic relationships among members of established 11 genera in the *Caliciviridae* family on the basis of complete VP1 amino acid sequences. Bootstrap support from 1000 replicates is indicated for values > 80%. The scale bar indicates the number of substitutions per site.

We also examined the complete coding genomic sequences of neboviruses for recombination events using RDP v4.101 (13). There was no recombination breakpoints found in any of the sequences. Furthermore, we optimized an in-house RT-PCR assay employing degenerate oligonucleotides targeting the capsid coding region (BoCalV_Cap_FP 5′-TACARAAGGAAGCACTC-3′, BoCalV_Cap_RP 5′-GGCGTACATGCGATTGAA-3′) to detect the presence of UK-B6-like caliciviruses in bovine calves in India (Appendix). Using this optimized RT-PCR, testing of 120 archived bovine diarrhoeic fecal samples, 40 each from the Indian states of Uttar Pradesh, Haryana and Himachal Pradesh, revealed frequent circulation of UK-B6-like caliciviruses in the Indian bovine population, with an overall positivity rate of 64.17% (77/120). In contrast, previous studies found nebovirus positivity rates of 12.0 % (6/50) in Turkey (2), and 5.2% (13/250) in Sweden (14).

## Conclusions

A bovine population of approximately 302.79 million have a significant impact on the livelihoods of farmers in India. Nonetheless, various enteric pathogens pose a significant threat to these animals’ health, resulting in substantial economic losses. During the current study, we identified a novel bovine calicivirus in an Indian diarrheic calf that is distantly related to *Nebovirus* and meets the ICTV criteria for designation of a probable new genus. Although it has been demonstrated experimentally that *Nebovirus* prototypes such as Newbury-1 and Nebraska can cause enteritis, further research is required to determine the enteritis-causing potential of UK-B6 calicivirus detected in India. Moreover, a broader investigation into the prevalence of UK-B6 type viruses and their potential co-infections or associations with other pathogens responsible for enteritis is required.

## Acknowledgments

The authors thank Dr. D.T. Maurya, Former Director of ICMR-NIV, Pune and Dr. R.K. Singh Former Director, ICAR-IVRI for the support.

