## Appendix Table 1; Appendix Table 2 for "A novel highly divergent enteric calicivirus in a bovine calf, India"

### Supplementary Information

**Appendix Table 1. Comparison of conserved motifs observed in Caliciviruses**

| **Proteins** | **Conserved motifs in Caliciviruses** | **Newbury agent 1** | **Nebraska virus strain NB** | **Mukti/India/UK-B6/2016** |
|---|---|---|---|---|
| 2C-like helicase-nucleoside triphosphatase DNA-binding | GXXGXGKS/T | GPPGHGKS (456-463) | GPPGHGKS (456-463) | GPPGVGKT (464-471) |
| ATP hydrolysis | KXXXFXSXXXXXS/TTN | KTRCFDSKYVLITTN (532-546) | KTRCFDSKYVIITTN (532-546) | KRTPFNSKFVIATTN (540-554) |
| 3D pol | GLPSG and YGDD | GLPSG (1440-1444), YGDD (1485-1488) | GLPSG (1440-1444), YGDD (1485-1488) | GLPSG (1451-1455) YGDD (1496-1499) |
| VP1 motif | PPG | PPG (1791-1793) | PPG (1791-1793) | PPG (1805-1807) |

**Appendix Table 2. VP1 amino acid sequence divergence of Indian Bovine Calicivirus (Mukti/India/UK-B6/2015) to all the representative members of the *Caliciviridae* family**

| **Genus** | **Caliciviruses** | **Divergence (%)** |
|---|---|---|
| Bavovirus | Calicivirus_chicken/Bavaria04V0021/DE/2004_HQ010042 | 80.9 |
| | Chicken_calicivirus_D62/2013_KM254170 | 80.9 |
| | Chicken_calicivirus_Q45/2013_KM254171 | 80.9 |
| Lagovirus | Rabbit_hemorrhagic_disease_virus_isolate_Sr12_2_KC741409 | 78.1 |
| | Rabbit_hemorrhagic_disease_virusFRG_M67473 | 78.7 |
| | Rabbit_calicivirus1_X96868 | 78.9 |
| | European_brown_hare_syndrome_virus_Z69620 | 77.8 |
| Minovirus | Fathead_minnow_calicivirusUSA/MN/2012_KX371097 | 90.9 |
| Nacovirus | Turkey_calicivirus_L11043_JQ347522 | 80.9 |
| | Chicken_calicivirus_F10026n_JQ347523 | 84.1 |
| | Goose_calicivirus_strain_N_KJ473715 | 83.3 |

| | | |
|---|---|---|
| Nebovirus | Newbury1_virus_DQ013304 | 65.8 |
| | Bovine_enteric_calicivirus_NB_AY082891 | 66.3 |
| | Bovine_calicivirus_Kirklareli_KT119483 | 9.4 |
| Norovirus | Lion_norovirus_GIV_2/Pistoia/387/06/ITA_EF450827 | 84.1 |
| | Swine_calicivirus_Sw918_AB074893 | 82.4 |
| | Bovine_norovirus_Newbury2_AF097917 | 82.4 |
| | Hu/NLV/Alphatron/982/1998/NET_AF195847 | 83.8 |
| | Bovine_calicivirus_Jena_AJ011099 | 83.8 |
| | Chiba_virus/GVIII_AJ844470 | 81.9 |
| | Murine_norovirus_1_AY228235 | 83.5 |
| | Sheep_norovirus_Norsewood_EU193658 | 82.9 |
| | Dog_norovirus_GVI1/HKU_Ca026F/2007/HKG_FJ692500 | 83.1 |
| | Dog_norovirus_GVI_1/Bari/91/2007/ITA_FJ875027 | 82.9 |
| | Dog_norovirus_Viseu_GQ443611 | 83.4 |
| | Rn/GV/HKU_CT2/HKG/2011_JX486101 | 82.7 |
| | Yuzawa_virus_GVIII_KJ196291 | 81.9 |
| | Bat_norovirusYN2010_KJ790198 | 80 |
| | Southampton_virus_L07418 | 82.6 |
| | Norwalk_virus_M87661 | 83 |
| | California_sea_lion_norovirus_strain_Csl/NoV2/PF0809162_MG572715 | 82.6 |
| | Hawaii_calicivirus_U07611 | 81.6 |
| | SapporoHK299_virus_GIX_1_KJ196290 | 81.7 |
| | Lordsdale_virus_X86557 | 81.2 |
| Recovirus | Tulane_virus_EU391643 | 86.7 |
| | Human_recovirus_Bangladesh_JQ745645 | 85.9 |
| | Human_Recovirus_Venezuela_MG571787 | 85.5 |
| | WUHARV_Calicivirus_1_JX627575 | 85.9 |
| | Tulane_Virus_FT205)_KC662363 | 85.5 |
| Salovirus | Atlantic_salmon_calicivirus_Nordland/2011_KJ577139 | 88.9 |
| | Atlantic_salmon_calicivirus_AL_V901_KJ577140 | 89.1 |
| Sapovirus | Porcine_enteric_calicivirus_Cowden_AF182760 | 77.3 |
| | Mex340_virus_AF435812 | 80.4 |
| | Houston_virus_71181_AF435814 | 79.1 |

| | | |
|---|---|---|
| | Arg39_virus_AY289803 | 79.7 |
| | NongKhai24_virus_AY646856 | 79.7 |
| | Porcine_sapovirus_JJ681_AY974192 | 82.7 |
| | Ehime_virus_DQ058829 | 77.7 |
| | Angelholm_virus_SW278_DQ125333 | 77.4 |
| | Porcine_sapovirus_2053P4_DQ359100 | 81.2 |
| | Porcine_sapovirus_43_EU221477 | 78.7 |
| | Porcine_sapovirus_sav1_FJ387164 | 77.8 |
| | Porcine_sapovirus_F1910_FJ498786 | 78.6 |
| | Sapovirus_MT2010/1982_HM002617 | 79.3 |
| | Sapporo_virus_U65427 | 79.3 |
| | Houston_virus/90_U95644 | 78.9 |
| | Manchester_virus_X86560 | 79 |
| | Bristol_virus_AJ249939 | 80.4 |
| | London_virus/29845_U95645 | 81.3 |
| Valovirus | Calicivirus_pig/AB90/CAN_FJ355928 | 84.1 |
| | Calicivirus_pig/F1510/CAN_FJ355929 | 84.3 |
| | Calicivirus_pig/AB104/CAN_FJ355930 | 83.9 |
| Vesivirus | Canine_calicivirusno48_AF053720 | 80.9 |
| | Pan1_virus_AF091736 | 82.5 |
| | Vesicular_exanthema_of_swine_virus_strain_A48_AF181082 | 82.5 |
| | Walrus_calicivirus_AF321298 | 81.9 |
| | Calicivirus_2117_AY343325 | 82.6 |
| | Canine_vesivirus_Bari/212/07/ITA_JN204722 | 82.7 |
| | Feline_calicivirus9_M86379 | 81.6 |
| | San_Miguel_sea_lion_virus1_M87481 | 82.1 |
| | San_Miguel_sea_lion_virus4_M87482 | 82.2 |
| | Feline_calicivirus_CFI/68_U13992 | 82.7 |
| | San_Miguel_sea_lion_virus17_U52005 | 82.4 |
